# A replication study on key neurodevelopmental pathways affected by valproate treatment of neural precursor cells

**DOI:** 10.1101/2022.11.21.517445

**Authors:** Salil K. Sukumaran, M Srividya, Shruti Iyer, Anushka Banerjee, Meera Purushottam, Sanjeev Jain, Biju Viswanath, Reeteka Sud

## Abstract

Adults with bipolar disorder or epileptic seizures are commonly prescribed sodium valproate. *In utero* exposure to this drug is linked to a multitude of defects in normal brain development, from neural tube defects to autism spectrum disorders. During the course of brain development, neural precursor cells (NPCs) give rise to neurons and glia, and therefore to understand the valproate-induced defects, it is crucial to understand its effect on NPCs. Two NPC lines, both derived from healthy individuals, were used for all experiments. Cells were treated with 0.7mM valproate for one week. Fresh media (+/− drug) was replenished every alternate day. RNA was extracted on day 7 of drug treatment, and transcriptomics performed. All experiments were performed in biological replicates. Genes that showed >1-fold difference (with FDR adjusted q-value ≤ 0.05) were considered differentially expressed. We further investigated the interacting partners of the differentially expressed genes using PINOT, as well as cellular pathways using DAVID. Our primary endpoint of analysis were genes that were differentially expressed (DEGs) with valproate treatment in both the NPC lines used. We found 21 such genes that were common in the two lines. PINOT revealed 504 interacting partners of the DEGs. Functional annotation analysis showed significant enrichment of four signaling pathways - Wnt, Notch, Rho-GTPase and PI3K-AKT. While the role of Rho-GTPase is a novel finding, we have replicated previously reported findings on Wnt, Notch and PI3K-Akt pathways, which further strengthens their role in mediating neurodevelopmental anomalies.

## 1. Introduction

Sodium valproate is a widely popular mood stabilizer, prescribed for patients with bipolar disorder (Cipriani et al., 2013). It is also commonly indicated for management of epilepsy; and though not as common, neuropathic pain, migraine, HIV polytherapies, and cancer (Lloyd and Sills, 2013). In pregnant women, the use of valproate can result in a wide variety of abnormalities in the developing fetus: cleft lip, limb defects, cardiovascular abnormalities, neural tube defects, developmental delays, etc.

Several aspects of valproate-induced teratogenicity have been shown over the years, including drug accumulation within the fetus, oxidative stress, folate antagonism and histone deacetylase inhibition, but the mechanisms linking these cellular processes to fetal malformations are still not clear despite intensive research. Reproducibility studies are therefore critically needed to ascertain the usefulness of the different mechanisms, and the contributions of different developmental pathways in the birth defects from valproate use. With this primary objective, we examined gene expression changes in human neural precursor cells (NPCs), derived from directed differentiation of induced pluripotent stem cells (IPSCs).

Neural stem cells (NSCs) are cells of the nervous system, which have the ability to self-renew, proliferate and differentiate into neuronal and glial lineages in the developing brain (Zhao and Moore, 2018). Human NPCs provide an opportunity to investigate human-specific aspects of development, disease mechanisms and drug-based studies. Exposure of NSCs to valproate was seen to promote neural differentiation while simultaneously inhibiting glial lineage specification and promote neurogenesis (Hsieh et al., 2004; Van Bergeijk et al., 2006; Chu et al., 2015). I*n vitro* treatment with valproate highlighted the roles of Wnt (Wang et al., 2015), Notch (Borghese et al., 2010), and Sonic hedgehog pathways (Zhang et al., 2020).

The present study validates valproate-induced modulation of these crucial neurodevelopmental pathways.

## 2. Materials and methods

### 2.1 Healthy control IPSCs

Two healthy control IPSC lines, C1 and C2, which were available in the Accelerator program for Discovery in Brain disorders using Stem cells (ADBS) biorepository (Viswanath et al., 2018), were used for the experiments. Control C1 participated in the study after written informed consent, which was approved by the ethics committee of the National Institute of Mental Health and Neurosciences, Bengaluru, India. C1 IPSC line was generated from blood lymphocytes as per published protocol (Iyer et al., 2018). C2 is a well-characterized integration-free IPSC line XCL1 (XCell Science), which is also part of the ADBS biorepository.

### 2.2 Generation of NPCs from human IPSCs

IPSCs were characterised for pluripotency markers OCT4 and SOX2 by immunostaining (Mukherjee et al., 2018). Using StemPro Accutase (Gibco), a well-characterized, high-quality IPSC culture was enzymatically dissociated, then grown in suspension until day 7 in Embryoid Body (EB) medium. [Knockout DMEM (Gibco), 20% KOSR (Gibco), 0.1 mM Non-Essential Amino Acids (Gibco), 2 mM Glutamax, 1% Penicillin-Streptomycin (Gibco), and 0.1 mM Betamercaptoethanol (Gibco)]. From day 7 to day 14, EB medium was replaced for Neural Induction Medium. [DMEM/F12 (Gibco), N2 supplement (Gibco), 8 ng/ml bFGF (Gibco), 1x Glutamax (Gibco), 1x Penicillin-Streptomycin (Gibco), 1x Non-essential Amino Acids (Gibco) and 2 μg/ml Heparin (Sigma)]. The EBs were plated on dishes coated with Matrigel (Corning) and allowed to develop neural rosettes. After manually passaging the neural rosettes, the tertiary rosettes were mechanically dissociated via pipetting and plated as an NPC monolayer. The medium was then replaced with Neural Expansion Medium [DMEM/F12 (Gibco), N2 supplement (Gibco), B27 supplement without Vitamin A (Gibco), 8 ng/ml bFGF (Gibco), 1x Glutamax (Gibco), 1x Penicillin-Streptomycin (Gibco), 1x Non-essential Amino Acids (Gibco) and 2 μg/ml Heparin (Sigma)]. Quantitative immunolabelling with Nestin and Pax6 revealed comparable NPC differentiation from IPSC lines (Paul et al., 2020).

### 2.3 Characterization of neural precursor cells

Cellular characterization of NPCs was performed by immunocytochemistry for SOX2 and nestin (Winiecka-Klimek et al., 2015). Details on the generation of NPCs and their cellular characterization, summarised above, have also been described in earlier publications from the ADBS consortium (Mukherjee et al., 2019; Paul et al., 2020).

### 2.4 Valproate treatment of neural precursor cultures

All experiments were carried out with cells at similar densities and passage numbers. The cells were seeded on a tissue culture treated surface coated with Matrigel at a density of 1.2×10^5^ cells/cm^2^. 12 hours post-seeding, sodium valproate (Sigma) was added to the media at a working concentration of 0.7mM for 7 days. Media change was performed every other day along with the required drug concentration. All cellular assays and isolation of RNA were performed on day 7. Total RNA was extracted using the Qiagen RNeasy Mini Kit (Cat. No: 74104). All experiments were performed in biological replicates.

### 2.5 Differential gene expression analysis in valproate treated cultures

RNA-sequencing was performed on the Illumina^®^ Hi-Seq platform. Genes showing >1-fold difference with FDR adjusted q-value <0.05 were considered differentially expressed in response to valproate treatment of neural precursors. The DEGs were then used as seeds for identifying protein-protein interaction (PPI) data using the Protein Interaction Network Online Tool (PINOT) (Version 1.1) (Tomkins et al., 2020). Interactions with a score of ≥ 2 (i.e., interaction is either detected by at least 2 methods (method score) or reported in at least 2 publications (publication score)) were filtered to obtain the list of interacting proteins. The DEGs and their interacting proteins were input into The Database for Annotation, Visualization and Integrated Discovery (DAVID) 2021 to identify the enriched functional pathways (Huang et al., 2009; Sherman et al., 2022).

## 3. Results

NPCs C1 and C2 were treated with 0.7mM valproate for 7 days, which falls within the range of values used in treatment paradigms in similar studies (Supp. table 1). RNA-seq was performed to obtain the transcriptome profile of C1 and C2 post-treatment with valproate.

### 3.1 Transcriptomic Analysis

Genes that showed >1-fold difference with FDR adjusted q-value ≤ 0.05 were considered differentially expressed. 23 differentially expressed genes (DEGs) were identified in C1, and 246 DEGs were identified in C2 (Supp. data 1 and 2). The DEGs from C1 and C2 were compared for common genes, where 21 genes were identified (Fig.1A). The log2 fold change values for the DEGs are depicted in Fig.1B.

**Figure 1.**
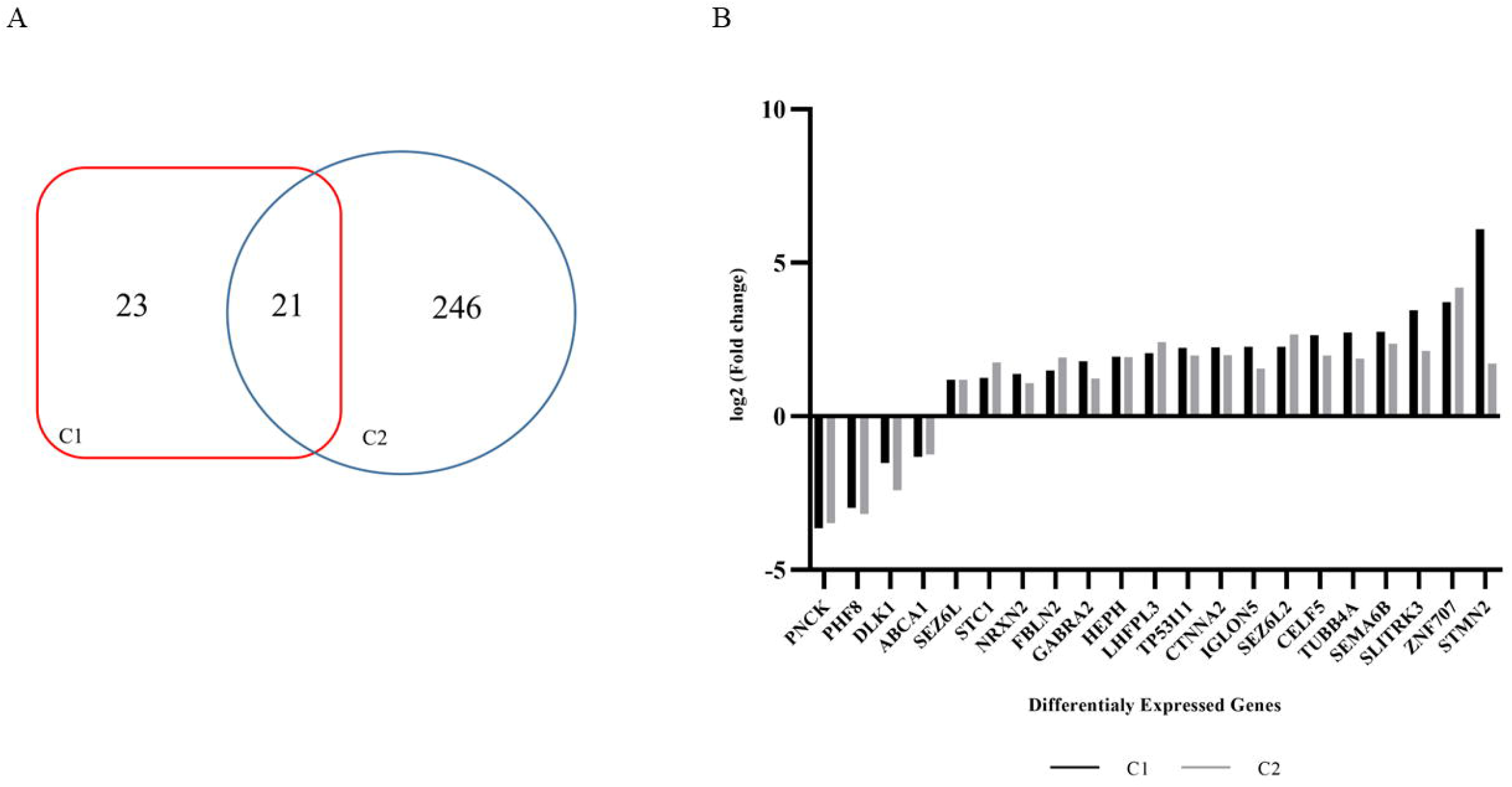
A: Venn diagram of common DEGs between C1 and C2 transcripts. Genes which showed >1-fold difference with FDR adjusted q-value < 0.05 were considered differentially expressed. 23 DEGs were identified in C1 and 246 DEGs were identified in C2, out of which 21 transcripts were found to be common between C1 and C2. B: Bar graph showing the log2 (Fold Change) for each of the 21 DEG transcripts for C1 (Black) and C2 (Grey).

Previously, Schulpen et al., (2015) exposed embryonic stem cell derived neural cultures to 1mM valproate for 7 days. The DEGs from the above study were compared with the 21 DEGs obtained in our analysis. 10 out of the 21 DEGs genes (*DLK1, ABCA1, NRXN2, FBLN2, CTNNA2, SEZ6L2, CELF5, TUBB4A, SLITRK3*, and *STMN2*) were differentially regulated in the same direction in Schulpen et al., 2015 (Table 1).

**Table 1:** Log2 fold changes values for each of the 21 DEG transcripts for C1 (Black) and C2 (Grey). 10 genes have been previously identified to be differentially expressed upon valproate treatment, while 11 genes have been identified in this study.

| Gene | C1 Fold change | C2 Fold change | Published evidence for role of valproate(1mM) in differential regulation |
| --- | --- | --- | --- |
| *PNCK* | -3.64473 | -3.48235 | No |
| *PHF8* | -2.98115 | -3.18254 | No |
| *DLK1* | -1.52233 | -2.40508 | Yes (Schulpen et al., 2015) |
| *ABCA1* | -1.31664 | -1.24286 | Yes (Schulpen et al., 2015) |
| *SEZ6L* | 1.19314 | 1.19096 | No |
| *STC1* | 1.26207 | 1.75739 | No |
| *NRXN2* | 1.37524 | 1.0788 | Yes (Schulpen et al., 2015) |
| *FBLN2* | 1.48833 | 1.91429 | Yes (Schulpen et al., 2015) |
| *GABRA2* | 1.79612 | 1.22786 | No |
| *HEPH* | 1.94731 | 1.93196 | No |
| *LHFPL3* | 2.05048 | 2.42193 | No |
| *TP53I11* | 2.22873 | 1.98249 | No |
| *CTNNA2* | 2.24829 | 1.9965 | Yes (Schulpen et al., 2015) |
| *IGLON5* | 2.26904 | 1.55205 | No |
| *SEZ6L2* | 2.27096 | 2.66991 | Yes (Schulpen et al., 2015) |
| *CELF5* | 2.64522 | 1.97755 | Yes (Schulpen et al., 2015) |
| *TUBB4A* | 2.72985 | 1.88081 | Yes (Schulpen et al., 2015) |
| *SEMA6B* | 2.75159 | 2.37348 | No |
| *SLITRK3* | 3.45606 | 2.125 | Yes (Schulpen et al., 2015) |
| *ZNF707* | 3.72458 | 4.19117 | No |
| *STMN2* | 6.09596 | 1.72267 | Yes (Schulpen et al., 2015) |

We report 11 additional genes (*PNCK, PHF8, SEZ6L, STC1, GABRA2, HEPH, LHFPL3, TP53I11, IGLON5, SEMA6B*, and *ZNF707*) that are found to be differentially expressed in NPCs upon valproate treatment (Table 1).

### 3.2 Earlier reports on valproate-regulated signaling pathways in different model systems

A literature survey was done to identify previous reports highlighting the role of valproate in modulating the enriched pathways (Table 2). Riva et al., 2018, reported that 2mM valproate treatment of glioma stem cells results in the upregulation of *WNT1* and its target genes. Greenblatt et al., 2008 showed that valproate treatment of medullary thyroid cancer cells resulted in an increase in both the full length and cleaved, active form of Notch1 and activates Notch1 signaling. A study by Lima et al., 2017 showed that valproate treatment in mouse model promotes antidepressant effects through increased hippocampal phospho-Akt (p-Akt) levels in WT, but not in PI3Kγ−/− mice. Pre-treatment with rapamycin, an inhibitor of mTOR, abolished the antidepressant-like effect of valproate, indicating the role of valproate in PI3K and mTOR activation (Lima et al., 2017). Another study by Zhang et al., 2017 also shows an increase in p-Akt in rat primary cortical neurons post-valproate treatment (Table 2). All of the above pathways are also reported to play an important role in neural tube closure defects (Rolo et al., 2018; Bekri et al., 2019; Han et al., 2020; Chen et al., 2021; Li et al., 2021; Tian et al., 2022).

**Table 2:**
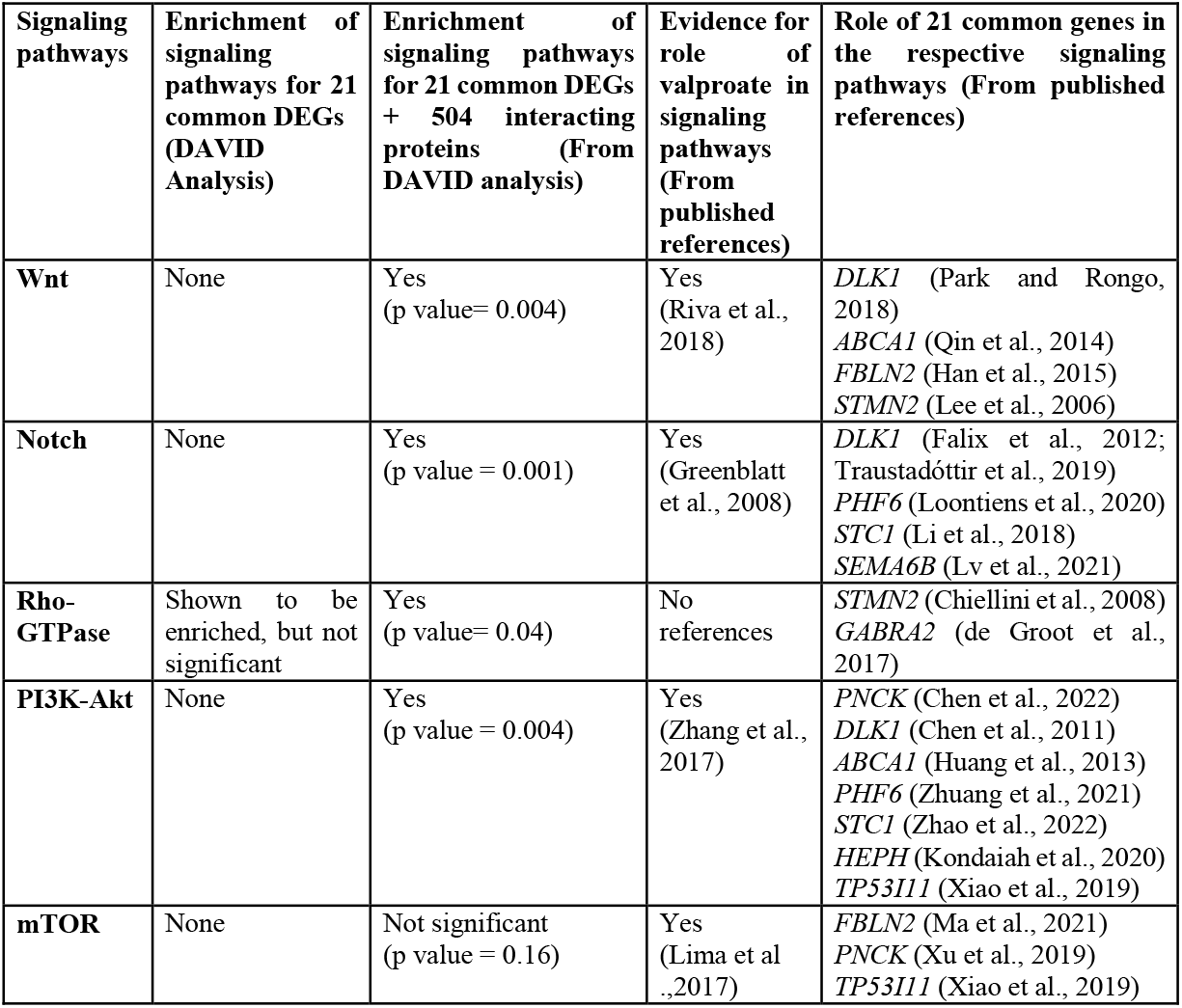
List of signaling pathways enriched for the DEGs and their interacting proteins, using DAVID analysis. Also shown in the table, previously published reports for the role of valproate in the enriched pathways.

### 3.3 Identification of functional protein interactions and pathways

The 21 common DEGs were analyzed using PINOT to determine protein-protein interactions (Supp. data 3). The first layer of interactions showed 504 interacting proteins for the DEGs. DAVID was used to identify the signaling pathways enriched by the 21 common genes, along with the 504 interacting proteins identified using PINOT (Supp. data 4). The analysis showed significant enrichment for Wnt, Notch, PI3K-Akt, and Rho-GTPase; however, mTOR pathway was not enriched in the analysis (Table 2). In addition, further literature review on the role of 21 common DEGs in the above-mentioned signaling pathways led to the identification of 11 DEGs that are involved in the pathways (Table 2).

## 4. Discussion

In observational studies, gestational exposure to valproate is associated with higher risks of cognitive, language, and psychomotor delay during early childhood and possibly with an increased risk of autism. Prospective study in pregnant women in developing countries showed an increase in congenital malformations to ~10%, up from 1-3% in the general population (Kochen et al., 2011). Considering that women of childbearing age make up a significant proportion of bipolar disorder patients (Macfarlane and Greenharlgh, 2018), addressing the safety concerns associated with their psychiatric treatment for their unborn child is paramount. And therefore, knowing the mechanisms associated with teratogenic effects of valproate can help address a challenging clinical issue, i.e. to manage the mental health needs of mother-to-be, while minimizing risks of teratogenicity.

Our analysis supports earlier reports on the role of Wnt, Notch, PI3K-Akt signaling pathways in different model systems subsequent to valproate treatment. Additionally, we report that Rho-GTPase pathway is directly modulated by valproate (Table 2). Given the previously reported role of this pathway in neural tube closure (Rolo et al., 2018), it follows that valproate should regulate this pathway in eliciting its teratogenic effects. Functional assays may further validate these findings on the role of Rho-GTPase pathway in the mechanistic action of valproate.

Earlier studies have reported transcriptomic changes subsequent to valproate treatment via induction of neurogenesis in swine model (Nikolian et al., 2018), neuronal differentiation in murine model (Zhang et al., 2017; Duan et al., 2019), and promotion of neurosphere formation in NSCs (Qi et al., 2022). Studies in rodent models showed that valproate reduces cellular growth and neural development (Barrett et al., 2017). In non-human primates, valproate was shown to exert its teratogenic effects by decreasing neuronal and glial markers and synaptic proteins which affect neurogenesis, leading to neural tube defects in the developing fetus (Zhao et al., 2019).

The paucity of systematic studies of risk/benefit analysis in women exposed to psychiatric medications during pregnancy is a grave cause for concern. Previous reports have drawn attention to the need for pre-pregnancy counseling in women with mental illness (Cook et al., 2010; Desai et al., 2012; Toh et al., 2013). Our specific objective here was to investigate neurodevelopmental pathways dysregulated by valproate exposure. It is important to note that risks of developmental anomalies are by no means restricted to valproate treatment (Viguera et al., 2002). All psychotropic medications can cross the placental barrier, and therefore can adversely affect a pregnancy in a myriad of ways. The specific question of valproate-induced developmental defects is thus integrally linked to broader issue of management of severe mental illnesses in pregnant women; and their side effects must be weighed against risks of untreated neurological/psychiatric condition in the expectant mother. Further research could also include longitudinal follow-ups, investigating the development of children exposed to medications during pregnancy and breastfeeding, so that women can be appropriately counseled when planning their pregnancy.

Though valproate is prescribed for a multitude of conditions, two most common reasons for prescription in pregnant women are epilepsy and bipolar disorder. Women with epilepsy reportedly have a higher rate of complications during pregnancy, and a higher risk of having a baby with birth defects due to anticonvulsant drugs such as valproate. Results of epidemiologic analysis of valproate use, made public by USFDA, also indicated cognitive impairments in children born to mothers who took valproate during pregnancy. Thus there are wide-ranging neurodevelopmental sequelae subsequent to exposure to this drug. The problem is further compounded by the fact that evidence on the mechanistic basis of neurodevelopmental anomalies due to valproate, in a relevant human model system, has been scant. And therefore, signaling pathways reported in our data that are affected by valproate can serve as important leads, targeting which could aid in better management of clinical risks to the developing fetus.

## Acknowledgments

The authors would like to thank Dr. Ravi Muddashetty and Dr. Dasaradhi Palakodeti for providing computational facilities for transcriptome analysis; Ms.Chitra B. and Mr. Mallappa M. for technical support. We are grateful to the participants and their family for their cooperation, as well as to clinicians and staff at NIMHANS for their assistance.

## Contribution to the field statement

Valproate is an anti-epileptic drug that when administered in early pregnancy can lead to developmental abnormalities in the fetus. The mechanistic action of the valproate in neurodevelopmental defects is not yet completely elucidated. Differential gene expression studies show the role of valproate in regulating Notch, Wnt, Shh and PI3K-Akt signaling pathways. Our transcriptomic analysis of the control NPC lines treated with valproate replicates the earlier available data on 10 gene transcripts, in addition to 11 novel genes which are identified in our study. Our data replicates the role of valproate on all of the above-mentioned signaling pathways, in addition to a novel finding of Rho-GTPase pathway. Additionally, the role of the Rho-GTPase in the process of neurulation in mice is already reported. This further strengthens the role of the valproate in mediating neurodevelopmental defects via the already reported pathways in addition to Rho-GTPase.

