## Supplementary table 1 for "A replication study on key neurodevelopmental pathways affected by valproate treatment of neural precursor cells"

**Supplementary Table 1: Valproate treatment paradigms used in earlier studies.**

| **Reference** | **Treatment Paradigm** | **Cell Type** | **Results** |
| --- | --- | --- | --- |
| Chanda et al., 2019 | 25µM to 3M valproate for 96 hours | Mouse neurons, human NSC | Caused chronic impairments in neurite development, dendritic morphology and functional properties of developing neurons, but not those of mature neurons.  Valproate-mediated inhibition of histone deacetylase (HDAC) and glycogen synthase kinase-3 (GSK-3) pathways, which caused transcriptional down-regulation of many genes including MARCKSL1 |
| Fukuchi et al., 2009 | 5mM valproate for 12h | Rat cortical neurons | Valproate-treatment markedly altered gene expression (up-regulated 726 genes, down-regulated 577 genes). Enhanced histone acetylation in promoters of upregulated genes |
| Fujiki et al., 2013 | 0.5 mM valproate, dose response treatment of 0.1, 0.5, 3, 20 mM, 3 days | NPCs, ES derived glutamatergic neurons | Proapoptotic effect on NPCs in dose-dependent manner.  NPCs treated with valproate died within 24. Dose-dependent (0.1–20 mM) enhancement of level of histone H3 acetylation in NPCs of ES cell-derived glutamatergic neurons.  Valproate did not influence apoptosis in ES derived glutamatergic neurons at any concentration (due to the inhibition of HDACs)  Neurons also increase histone acetylation, but not as rapidly in NPCs. |
| Qi et al., 2022 | 1, 50, 100, 500, 1000, and 10 000 µmol/L for 48h | Rat NSCs | Enhanced neurosphere formation and NSC proliferation, upregulation of TGFB1 |
| Hsieh et al., 2004 | 0.3mM to 1mM valproate for 4 days, Adult female fisher rats- two daily injections of 300mg/kg valproate. | Rat NPCs | Induced neuronal differentiation of NPCs, inhibited astro and oligo differentiation. Upregulation of NeuroD transcription factor. |
| Jung et al., 2008 | 1mM valproate for 48 hours | Rat NPCs | Induces differentiation, reduces proliferation via ERK-p21^Cip/WAF1^ pathway. |
| Shinde et al., 2016 | 1mM valproate for 4 days | H9 hESCs, foreskin hiPSCs, IMR90 hiPSCs | Upregulation of PRKCB gene post treatment, suppression of upregulated developmental genes and induction of downregulated ones |
| Lu et al., 2017 | 1mM valproate upto 7 days | Mouse eNSCs | Promotes the neuronal differentiation of eNSCs and neurite outgrowth of NSC-derived neurons. Induces phosphorylation of c-Jun by JNK |
| Go et al., 2012 | 400mg/kg (pregnant Sprague-Dawley rats on E12), 0.2 mM (12hr), 0.5 mM (24h) | Rat NPCs | Macroencephaly in prenatal brain post exposure, increase in NPC proliferation |
| Fuller et al., 2002 | 0.75 - 3mM valproate for 24-48h | Chick embryo neural crest cells | Decreased individual migration of cells and cellular spreading. Valproate at 3mM arrested proliferation |
| Zhou et al., 2011 | 1mM valproate for 48h | Mouse NSCs | Blocks neurosphere formation, inhibited DNA synthesis |
| Wang et al., 2015 | 0.75mM valproate for 3, 7 and 10 days | Rat NSCs | Valproate induces neuronal differentiation of NSCs by activating Wnt signal pathway |
| Foti et al., 2013 | In vivo paradigm - 250ug/g on PN7 and 1h before culture on PN8  In vitro paradigm - 10mM valproate on day 0 and 4 days later | CD-1 mice | Inhibits NSC activity in vivo and in neurosphere expansion |
| Schulpen et al., 2015 | 0.1mM, 0.33mM and 1mM for 7 days | hESCs neural differentiation | The effect of valproate on differentially expressed genes during hESC differentiation were assessed |
